# ILC2s navigate tissue redistribution during infection using stage-specific S1P receptors

**DOI:** 10.1101/2024.05.12.592576

**Authors:** Takamasa Ito, Yoshihiro Ishida, Yingyu Zhang, Vincent Guichard, Wanwei Zhang, Richard Han, Kevin Guckian, Jerold Chun, Jianwen Que, Allen Smith, Joseph F. Urban, Yuefeng Huang

## Abstract

Lymphocytes can circulate as well as take residence within tissues. While the mechanisms by which circulating populations are recruited to infection sites have been extensively characterized, the molecular basis for the recirculation of tissue-resident cells is less understood. Here, we show that helminth infection- or IL-25-induced redistribution of intestinal group 2 innate lymphoid cells (ILC2s) requires access to the lymphatic vessel network. Although the secondary lymphoid structure is an essential signal hub for adaptive lymphocyte differentiation and dispatch, it is redundant for ILC2 migration and effector function. Upon IL-25 stimulation, a dramatic change in epigenetic landscape occurs in intestinal ILC2s, leading to the expression of sphingosine-1-phosphate receptors (S1PRs). Among the various S1PRs, we found that S1PR5 is critical for ILC2 exit from intestinal tissue to lymph. By contrast, S1PR1 plays a dominant role in ILC2 egress from mesenteric lymph nodes to blood circulation and then to distal tissues including the lung where the redistributed ILC2s contribute to tissue repair. The requirement of two S1PRs for ILC2 migration is largely due to the dynamic expression of the tissue-retention marker CD69, which mediates S1PR1 internalization. Thus, our study demonstrates a stage-specific requirement of different S1P receptors for ILC2 redistribution during infection. We therefore propose a fundamental paradigm that innate and adaptive lymphocytes utilize a shared vascular network frame and specialized navigation cues for migration.

## Introduction

Innate lymphoid cells (ILCs) mirror T helper subsets in terms of essential transcription factors, effector cytokines and immunological functions, while these two types of cells exhibit distinct responsive patterns in an event of infection(*1*). The cardinal feature of adaptive T cells is the antigen-dependent activation of naïve cells in secondary lymphoid sites, followed by the migration of effector cells through efferent lymph to the blood circulation and then to a peripheral tissue site of infection(*2*). Memory cells are generated during this process and continue patrolling the body for immunosurveillance, and a fraction of them take residence and become tissue-resident memory T (T_RM_) cells for localized protection(*3*). By contrast, ILCs are largely tissue-resident cells, adapted to their specific environments during development and performing effector functions locally upon cytokine stimulation(*4*). However, this view has been extended by the characterization of interorgan trafficking of ILC2s during helminth infection(*5*). In response to the alarmin signal IL-25, intestinal ILC2s undergo robust proliferation and subsequently enter circulation. These migratory ILC2s, termed inflammatory or induced ILC2s (iILC2s), are recruited to distal organs including the lung where they contribute to anti-helminth responses, particularly in the early stage of infection before T effectors are generated, as wells as in the circumstance of T cell deficiency(*6, 7*). The mechanisms of T cells egress from lymph nodes (LNs) and recruitment to infection sites have been extensively characterized, whereas the migratory route and molecular basis of intestine-resident ILC2 redistribution are poorly understood.

The tissue-draining lymphatic system is considered as the primary route for leukocytes exit from peripheral tissues and return to circulation(*8*). Extravasated fluid, macromolecules and leukocytes, for example, antigen-presenting dendric cells, are taken up by blind-ended lymphatic capillaries, which converge into larger lymphatic vessels that drain into regional lymph nodes (LNs). Upon passage through a chain of LNs, which are connected by adjoining collecting lymphatic vessels, lymph is then returned to the blood vasculature through the thoracic ducts, which merge into the subclavian vein(*9*). On the other hand, circulating immune cells in the bloodstream traverse the endothelial layer of blood vessels via a process known as transendothelial migration (TEM), to enter tissue sites of infection(*10*). TEM has been long thought as a unidirectional trip from blood vessel lumen to abluminal tissue, but increasing evidence supports that reverse TEM also occurs. For example, neutrophils directly return to blood circulation during tissue injury, suggesting the existence of an alternative route for leukocytes exit from tissue especially under pathological conditions(*11–14*). iILC2s were observed to be present within lymphatic vessels of the intestine(*7*), but whether they use lymphatic system as primary migration route during helminth infection is to be determined.

Immune cell migration is orchestrated by various molecules including selectins, chemokine receptors, integrins and chemoattractants(*15*). One of the pivotal determinants is sphingosine 1-phosphate (S1P), a lipid mediator that is enriched in blood and lymph while maintained low in interstitial fluids, creating a steep S1P gradient, which can be sensed by S1P receptor (S1PR)-expressing cells to facilitate the egress from lymphoid organs(*16–18*). For examples, S1PR1 is essential for T cell egress from thymus and LNs, and its downregulation in inflamed tissue contributes to forming T_RM_ cells(*19–21*). NK cell egressing from bone marrow and LNs predominantly relies on S1PR5(*22, 23*). The migration of activated dendritic cells from tissues into regional LNs depends on S1PR1 and S1PR3(*24–26*). S1PR1 and S1PR4 are also reported to regulate effector T cells recirculation from peripheral tissue back to the draining LNs(*27*). We previously showed that migratory iILC2s upregulate the expression of a few S1PRs and FTY720 treatment, a pan-S1PR antagonist(*28, 29*), blocks iILC2 accumulation in periphery tissues. However, which S1PR(s) play a major role during ILC2 migration and which step(s) of the migration they regulate are unknown.

Here, we characterized a migratory path of intestinal ILC2s recirculation and revealed a stage-specific requirement of different S1P receptors for ILC2 trafficking. Upon helminth infection, intestinal ILC2s enter local lymphatic capillaries to access lymph-to-blood circulation but the secondary lymphoid structure is not required for ILC2 migration and effector function. Unlike Th2 gene loci, which acquire accessibility in ILC2s during development and change little after activation, chromatin regions in proximity to S1P receptor genes are inaccessible in intestinal ILC2s at the steady state but undergo a dramatic change in response to IL-25 stimulation, enabling the expression of S1PR1 and S1PR5. While S1PR5 regulates ILC2 exit from intestinal tissue to lymph, S1PR1 predominately modulates ILC2 egress from mesenteric LNs. The tissue retention marker CD69 negatively regulates the presence of S1PR1 on ILC2 cell surface, contributing to the basis of the stage-specific requirement of different S1PRs for ILC2 redistribution during infection. These findings suggest a fundamental paradigm that ILCs and T cells utilize a shared vascular network frame and specialized navigation cues for migration.

## Results

### ILC2 redistribution from gut to periphery during infection requires the access to the tissue-draining lymphatic system

To determine whether intestinal ILC2s access circulation by travelling through lymphatic vessel network, we sought an approach to cut off lymph flow from the gut to peripheral circulation. Gut-draining mesenteric LNs (MLNs) are the junction points of collecting lymphatic vessels from intestinal tissue and are visibly identifiable in mice (Figure 1A). We surgically excised MLNs in wild-type C57BL/6 mice, to cut off the access of local lymphatic vessels to the main lymph (Figure 1B). Three-day intraperitoneal injection of IL-25 cytokine elicited gut-derived KLRG1^hi^ ST2^−^ iILC2s in the lungs and peripheral blood but failed to do so in MLNs-removed mice (Figure 1C-E), suggesting that the migration of gut-derived iILC2s to periphery requires the access to lymphatic vessel network.

**Figure 1.**
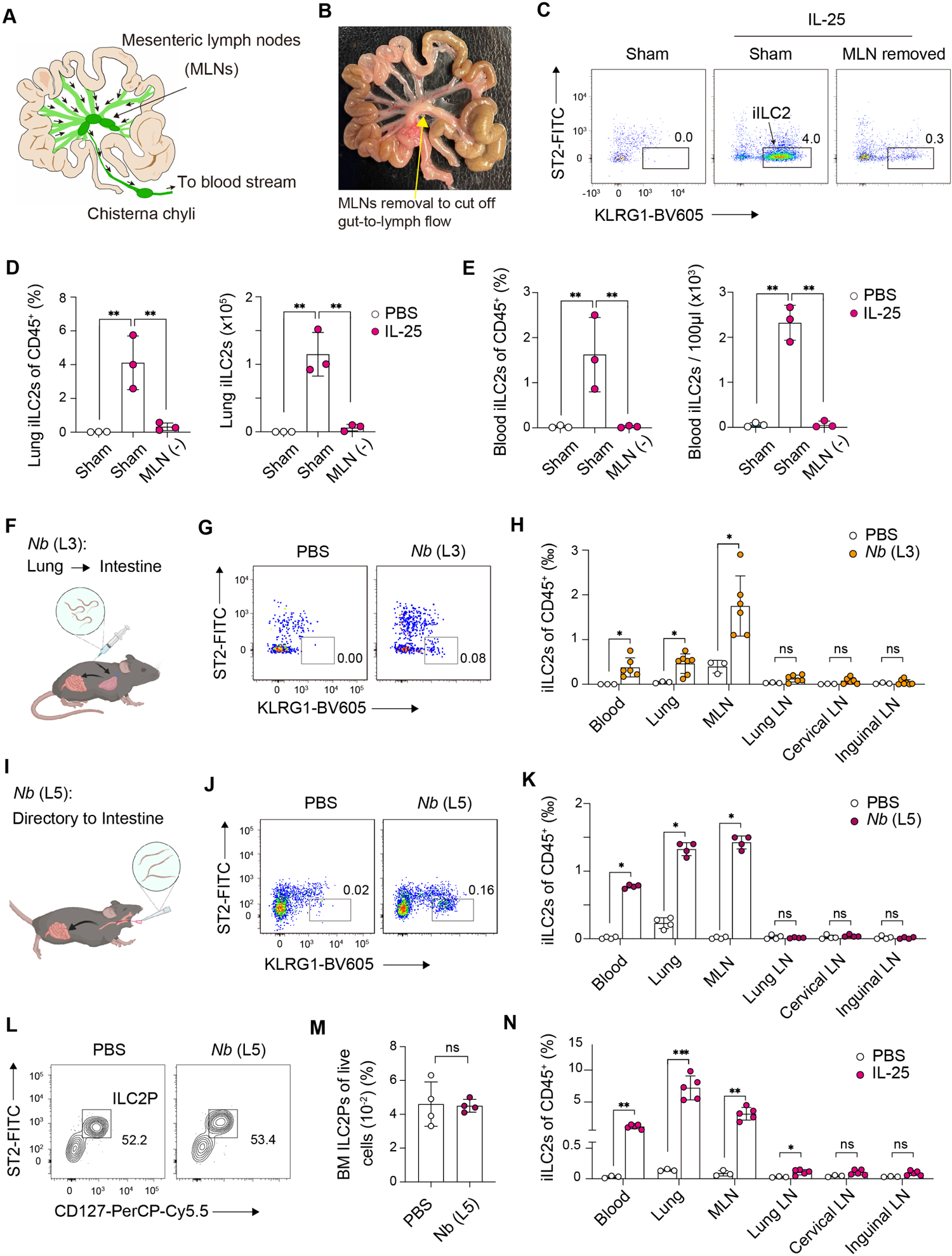
Intestinal ILC2 redistribution requires the access to the tissue-draining lymphatic system. **(A** and **B)** A picture and diagram of mesenteric lymph nodes (MLNs). Black arrows in **A** shows the direction of the lymph flow. **(C-E)** Flow cytometry (FACS) analysis of iILC2s in the lung (C and D) and in the blood (E) of sham or MLN-excised C57BL/6 mice, which were i.p. injected with IL-25 for three days. iILC2s were gated as live CD45^+^ Lin^−^ Thy1^+^ GATA3^+^ KLRG1^+^ ST2^−^. **(F)** Schematic of *N. brasiliensis* (*Nb*) L3 infection route when L3 larvae were subcutaneously injected into the mice. **(G)** FACS analysis of iILC2s in the lung of B6 mice on day 5 post subcutaneous injection of 350 *Nb* L3. **(H)** FACS analysis of the frequency of iILC2s in individual organs of the mice in G. **(I)** Schematic of *Nb* L5 infection route when L5 larvae were orally gavaged to the mice. **(J)** FACS analysis of iILC2s in the lung of B6 mice on day 4 post oral gavage of 350 *Nb* L5. **(K)** Frequency of iILC2s in individual organs of the mice in J. **(L, M)** FACS analysis of ILC2 progenitor (ILC2P) in the bone marrow (BM) of the mice in J. **(N)** Frequency of iILC2s in individual organs of the mice receiving three-day i.p. injection of IL-25. Data in (D), (E), (H), (K), (M), (N) are shown as the mean ± SEM. Student *t* test; ns, not significant; *p < 0.05, **p < 0.01, ***p < 0.001. Data are representative of two independent experiments.

We then tested the possibility that whether intestinal ILC2s can directly enter blood vessels via reverse TEM. If cells entered bloodstream firstly, they would be systemically distributed into many peripheral LNs through high endothelial venules (HEVs). Thus, we surveyed the presence of iILC2s in LNs at different tissue sites during *Nippostrongylus brasiliensis* infection. On day 5 post inoculation of *N. brasiliensis* stage 3 larvae (L3), iILC2s were detected in blood, lung and MLNs, but not in mediastinal, cervical or inguinal LNs (Figure 1F-H), suggesting that intestinal ILC2s don’t directly access bloodstream; instead, they use tissue-draining lymphatic network as their primary route for tissue exit during helminth infection.

In mice, *N. brasiliensis* L3 migrate through the blood and tissues to the lung, then are coughed up and swallowed, and subsequently reach the small intestine, where they complete the rest of life cycle(*30*). It has been suggested that bone marrow (BM) progenitor cells also contribute to circulating ILC2 populations during *N. brasiliensis* infection(*31*). The mobilization of lung-resident ILC2s during allergic inflammation was also reported (*32*). Thus, we employed a model of oral gavage of *N. brasiliensis* stage 5 larvae (L5), which directly go to gastrointestinal tract without damaging lung and other tissues(*33*) (Figure 1I). Similar to *N. brasiliensis* L3, L5 infection elicited iILC2s in blood, lung, MLNs but not in other peripheral LNs (Figure 1J, K), and did not induce BM ILC2 progenitor expansion (Figure 1L, M), further supporting that migratory iILC2s are largely derived from the intestine. Three-day IL-25 injection mimicked *N. brasiliensis* infection in the redistribution of intestinal ILC2s (Figure 1N), so we used IL-25 treatment as a model to further investigate the mechanism of ILC2 migration.

### Secondary lymph structures are redundant for ILC2 migration and effector function

To illustrate the migratory path of iILC2s in gut-draining lymphatic system, we performed confocal imaging of MLNs in IL-25-treated mice. The majority of iILC2s were localized in T cell zone and subcapsular sinus regions, where afferent lymphatic vessels merge into MLNs, but very few in B cell zone (Figure 2A-C), suggesting that the majority of iILC2s travel through MLNs rather than take a residency (Figure 2D).

**Figure 2.**
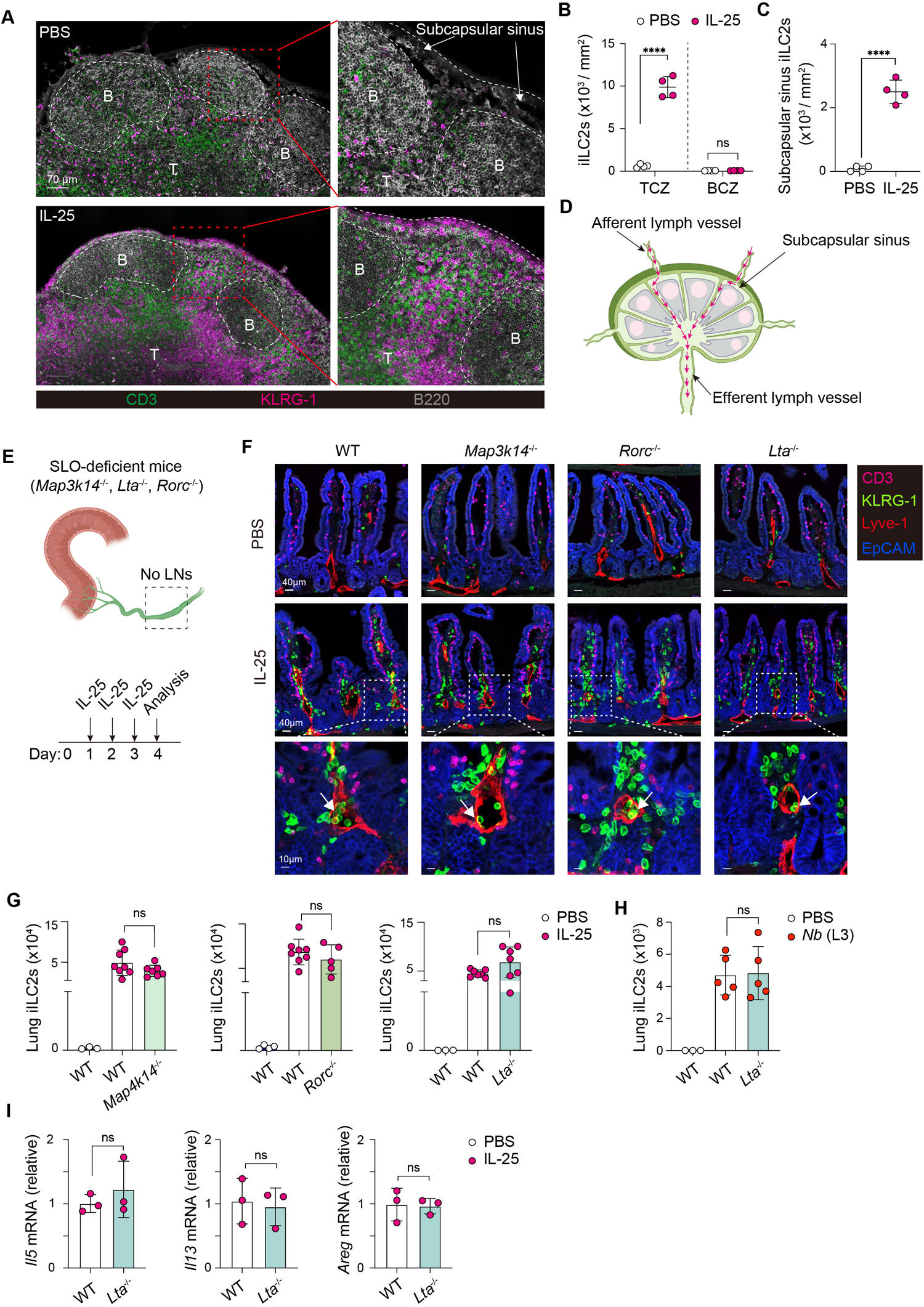
Secondary lymph structures are redundant for intestinal ILC2 redistribution. **(A)** Confocal imaging of MLNs from B6 mice treated with or without IL-25. T, T cell zone; B, B cell zone. **(B** and **C)** Quantitative analysis of iILC2 in T cell zone (TCZ), B cell zone (BCZ) and subcapsular sinus in A. **(D)** Schematic of iILC2 migratory path through MLNs (red arrows). **(E)** Experimental schematic of iILC2 induction in three secondary lymph organ (SLO) deficient mouse lines. **(F)** Confocal imaging of small intestine from the mice treated with PBS or IL-25 as in E. **(G)** FACS quantification of iILC2s in the lungs of the mice in F. **(H)** iILC2 cell numbers in the lung of the mice infected with *Nb* L3 for 5 days. **(I)** RT-qPCR analysis of *Il5*, *Il13*, *Areg* mRNA expression in the lung of the mice treated with IL-25 for 3 days. Data in (B), (C), (G), (H), (I) are shown as the mean ± SEM. Student *t* test; ns, not significant; *p < 0.05, **p < 0.01, ***p < 0.001, ****p < 0.0001.

Lymph nodes and other secondary lymphoid organs (SLOs) evolute along with adaptive lymphocytes, to provide a signal hub where a large pool of circulating naïve T cells scan a rare antigen presenting cell for clonal expansion (*34, 35*). We tested whether the secondary lymphoid structure is required for the migration and effector function of innate ILC2s, by using SLO-deficient mouse strains including *Lta*^−/−^, *Rorc*^−/−^ and *Map3k14*^−/−^(*36–38*) (Figure 2E and F). Using different genetic strains with the similar SLO-deficient phenotype would exclude the cell-intrinsic effect of individual gene deletion on ILC2 *per se*. There was no difference in cell numbers of intestinal ILC2s between WT and SLO-deficient mice at steady state or after IL-25 treatment (Supplemental Figure 2), suggesting that SLO deficiency doesn’t impact ILC2 development or cytokine-induced *in situ* proliferation. Imaging of the small intestine tissue showed that *Lta*^−/−^, *Rorc*^−/−^ and *Map3k14*^−/−^ mice had normal lymphatic capillaries, and IL-25-stimulated KLRG1^+/hi^ ILC2s were able to enter lymphatic vessels in these SLO-deficient mice, similar to WT mice (Figure 2F). The cell numbers of IL-25- or *N. brasiliensis* infection-elicited iILC2s in the lungs of SLO-deficient mice were similar to the numbers in WT mice (Figure 2G and H). The mRNA levels of *Il13*, *Il5* and amphiregulin (*Areg*) in iILC2s from SLO-deficient mice were similar to WT mice (Figure 2I). These results suggest that lymph nodes and other SLOs are redundant for the migration and effector cytokine production of iILC2s.

### IL-25 induces a dramatic epigenome remodeling in ILC2s to enable the expression of S1PRs

We next investigated the molecular basis of ILC2 migration. We firstly performed bulk RNA sequencing (RNA-Seq) of steady-state gut ILC2s (gILC2s) and IL-25-induced migratory iILC2s at different tissue sites including gut (yet to egress), mLNs and lung. Principal component analysis (PCA) demonstrated that the transcriptome profile of gut iILC2s was placed between resident gILC2s and mLN/lung iILC2s (Figure 3A), suggesting that gut iILC2s represent a transition phase from steady-state resident gILC2s to migratory iILC2s in periphery. Heatmap analysis confirmed that, while the gene expression patterns of MLN and lung iILC2s were significantly distinct from gILC2s, gut iILC2s showed the partial resemblance to both groups (Figure S3A). Gene sets related to chemokine-receptor binding, innate immune response and intestinal absorption were highly enriched in MLN iILC2s compared with gILC2s (Figure 3B and S3C), indicating the active and migratory feature of iILC2s. Among many chemoattractant receptors, the expression of S1PRs was dramatically increased in migratory iILC2s compared with resident gILC2s, whereas CCR7, which is important for the trafficking of DCs and ILC3s between gut and mLNs, was not upregulated in iILC2s (Figure S3B), suggesting that ILC2s undergo a specific transcriptomic reforming for their migration.

**Figure 3.**
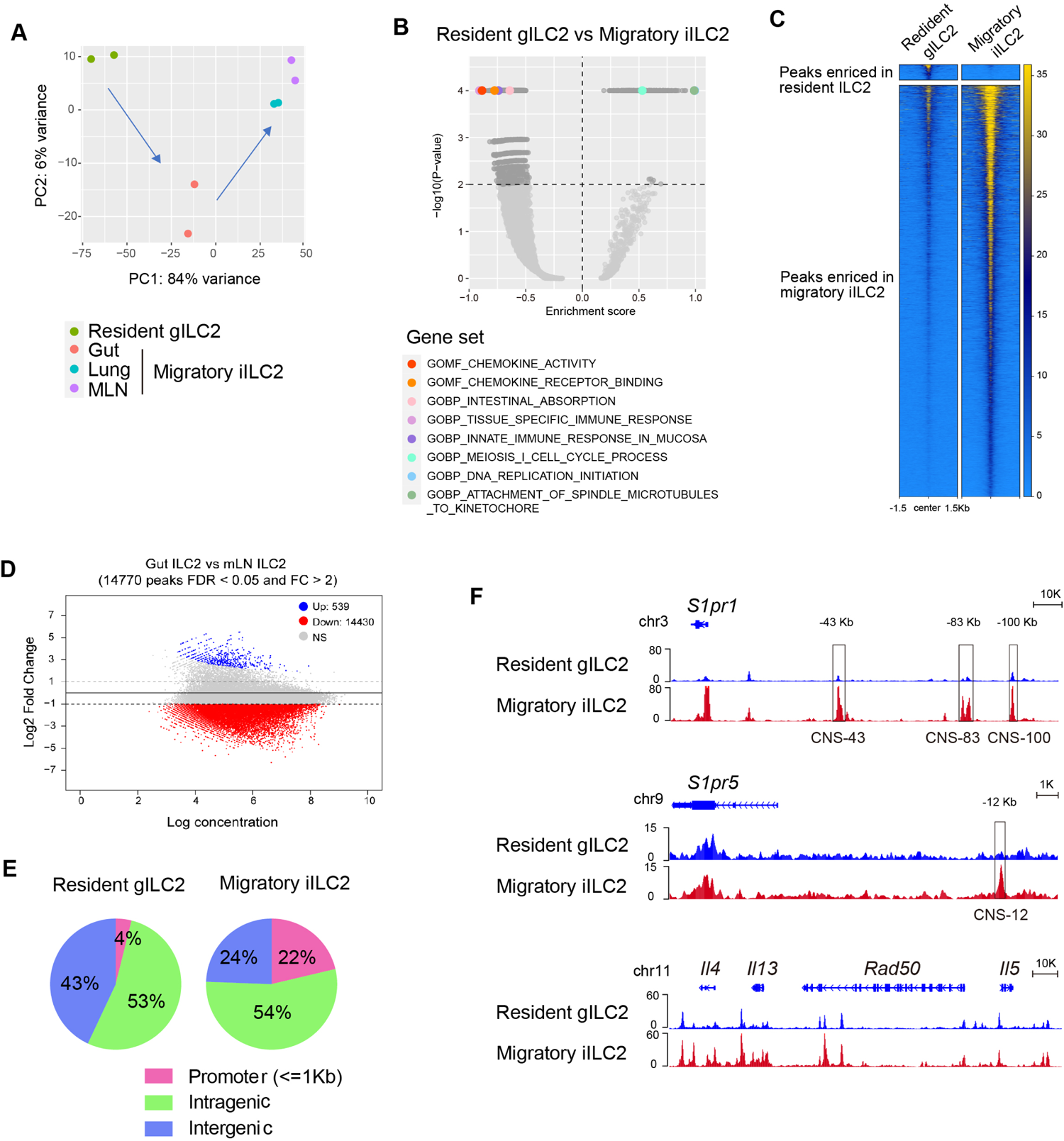
IL-25 induces a selective epigenome remodeling in migratory ILC2s. **(A)** Principal components analysis (PCA) of bulk RNA-Seq of sorted intestinal ILC2s from untreated B6 mice (resident gILC2) and sorted iILC2s from small intestine, MLNs and lung of IL-25-treated B6 mice. **(B)** Volcano plot of gene sets enriched in resident gILC2s and MLN iILC2s, respectively. **(C)** Heatmap of the distribution of ATAC peaks around start sites ±1.5kb. ATAC-Seq was performed using sorted gILC2s and MLN iILC2s as in A. **(D)** MA Plot of ATAC peaks enriched in resident gILC2s and migratory MLN iILC2s. The number of peaks differential identified with a FDR<0.05 and FC>2. **(E)** Pie charts illustrate the distribution of ATAC peaks across the genome including promoter ±1kb of transcription start sites, intragenic and intergenic regions. **(F)** Genomic track view of accessible ATAC peaks in proximity to *S1pr1*, *S1pr5*, and Th2 cytokines loci in gILC2s and MLN iILC2s.

To understand how the upregulation of migration-associated genes in ILC2s is modulated, we performed an assay for transposase-accessible chromatin with sequencing (ATAC-Seq) of resident gILC2 and IL-25-elicited MLN iILC2s. We found that three-day IL-25 injection induced a dramatic, genome-wide increase in chromatin accessibility (Figure 3C). 14,770 peaks were found highly accessible in migratory iILC2s but not in resident gILC2s, while only 539 peaks were highly accessible in gILC2s but not in iILC2s (Figure 3D). Analyzing the genomic distribution of ATAC peaks revealed that the percentage of accessible promoter regions within ±1kb of transcription start sites (TSS) were significantly higher in iILC2s (22%) compared that in gILC2 (4%) (Figure 3E), indicating that the changes of chromatin landscapes caused by IL-25 stimulation appear to be primarily shaped by promoter regions. Within the *S1pr1* locus, multiple regulatory element (RE) regions gained great accessibility in iILC2 compared to gILC2, including CNS-43 (conserved non-coding sequence located 43 kilobases upstream of the TSS), CNS-83, and CNS-100 (Figure 3F). Similarly, a peak at CNS-12 region of *S1pr1* locus appeared in iILC2 compared to gILC2. By contrast, the accessibility in type 2 effecter gene loci remained largely unchanged in iILC2 compared to gILC2 (Figure 3F), consistent with the previous report that ILC2s acquire chromatin accessibility in proximity to Th2 effector genes during the development and change little after activation(*39*). These results suggest that IL-25 induces a selective epigenome remodeling to enable the expression of S1PRs for ILC2 migration.

We then investigated the mechanisms of transcriptional control of S1PRs expression. Motif searching of transcription factor (TF) binding sites within ATAC peaks in iILC2s revealed top five TF candidates including RUNX1, ETS1, KLF2, BATF and IRF3 (Figure S4A), suggesting that these TFs potentially regulate transcriptional program in iILC2s. Among them, KLF2 has been reported to be a key TF to regulate S1PR1 expression(*40*). FIMO analysis showed that all three accessible REs in *S1pr1* locus contained KLF2 binding sites (Figure S4B), suggesting a possible role of KLF2 in upregulating S1PR1 in iILC2s. In *S1pr5* locus, the accessible RE region contained the binding sites for ZEB2 (Figure S4C), a TF that has been showed to modulate S1PR5 expression in CD8+ T cells (*41*). The TE regions in proximal to *Klf2* and *Zeb2* genes also acquired accessibility in iILC2s compared to gILC2s (Figure S4D and E), and these accessible RE regions contained binding sites for BATF (Figure S4F and G), a TF that plays an important in the induction of iILC2(*42*). Consistently, BATF transcription activity was significantly increased in both MLN iILC2s and lung iILC2s compared to gILC2s (Figure S4H). Together, these results suggest that BATF putatively orchestrates the transcriptional regulation cascade to mediate the expression of S1PRs in ILC2s.

### S1PR1 and S1PR5 regulate ILC2 migration both *in vivo* and *in vitro*

To test whether S1P-mediated chemotaxis is critical for ILC2 migration, we used a S1P-lyase inhibitor 4-deoxypyridoxine (DOP), which blocks irreversible degradation of S1P and leads to S1P gradient disruption(*43*). DOP treatment inhibited IL-25-induced iILC2 redistribution to the lung (Figure 4A and B), suggesting that a S1P gradient is required for ILC2 migration. Notably, DOP treatment resulted in a decrease in T cell number in the lung (Figure 4B), presumably due to T cell sequestration in lymphoid organs. We previously reported that FTY720 blocks ILC2 migration from gut to the lung(*7*). To determine which S1PR(s) are required for ILC2 migration, we employed different inhibitors that specifically target individual S1PR, including SEW2871 (for S1PR1), CYM50358 (for S1PR4) and BIO-027223 (for S1PR5) (*44–46*). SEW2871 or BIO-027223 treatment reduced the cell numbers of iILC2s in periphery blood and the lung of IL-25-treated mice in a dose-dependent manner (Figure 4C and E), indicating that S1PR1 and S1PR5 are important for ILC2 migration. CYM50358 treatment did not affect iILC2 cell numbers in the lung and blood (Figure 4D), suggesting that S1PR4 is dispensable for ILC2 redistribution. We sorted migratory iILC2s from IL-25-treated mice and performed transwell assay. Both S1PR1 and S1PR5 inhibitors suppressed S1P-mediated ILC2 transmigration in a dose-dependent manner (Figure 4F and G), suggesting that S1PRs directly modulate ILC2 migration. Thus, the upregulation of S1PR1 and S1PR5 is required for ILC2 migration.

**Figure 4.**
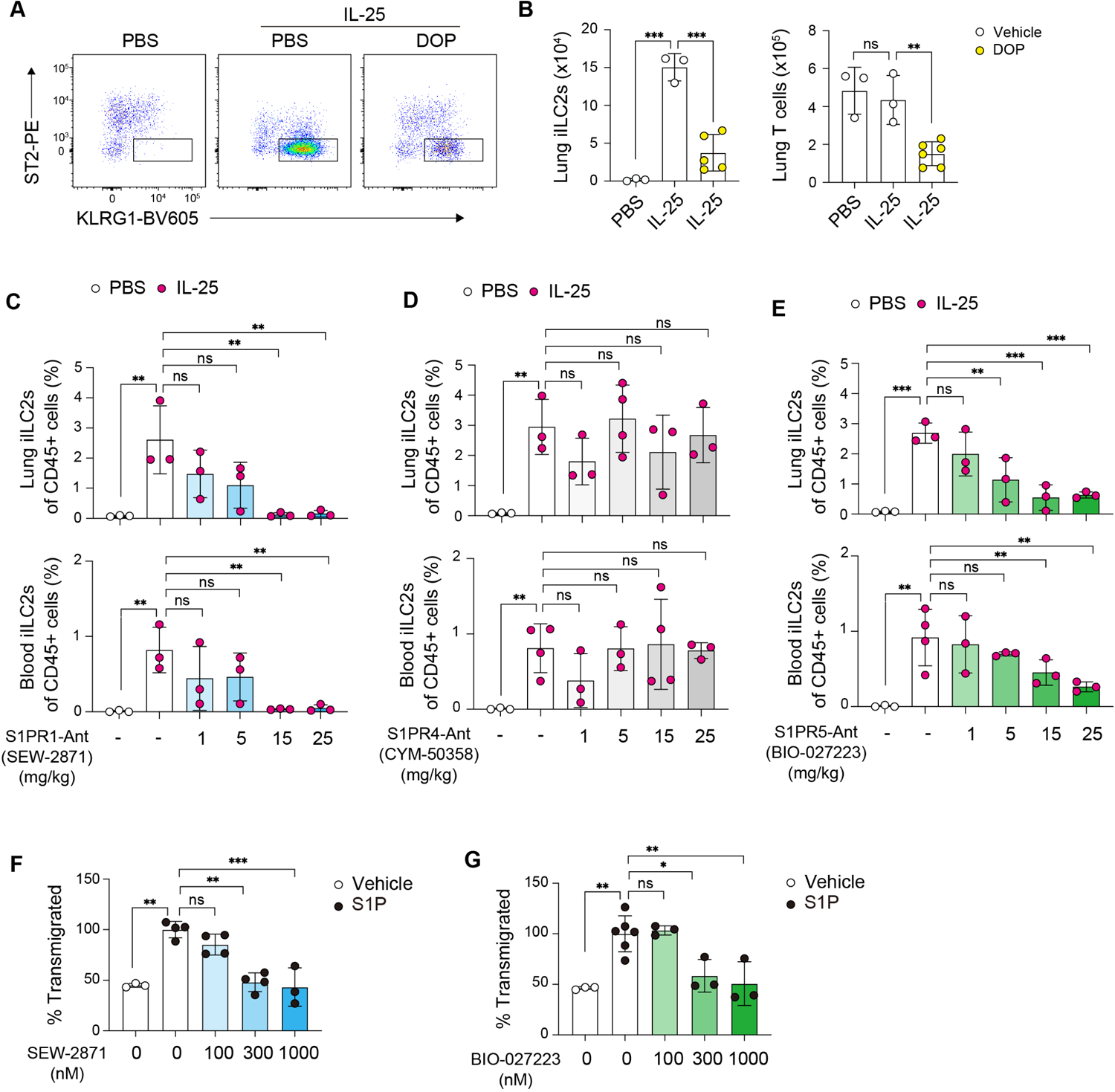
S1PR1 and S1PR5 regulate ILC2 migration both *in vivo* and *in vitro*. **(A)** FACS analysis of lung iILC2s in B6 mice receiving three-day IL-25 injection with or without 4-deoxypyridoxine (DOP) treatment. **(B)** Frequency of iILC2s and CD4+ T cells in the lung of the mice in A. **(C-E)** FACS quantification of iILC2 frequency in the lung and blood of B6 mice treated with IL-25 plus SEW-2871 **(B)**, CYM-50358 **(C)** or BIO-027223 **(D)** at the indicated doses. **(E** and **F)** Frequency of transmigrated iILC2s in transwell assays toward S1P. Cells were pretreated with or without SEW-2871 or CYM-50358 at indicated concentrations. Data is shown as the mean ± SEM. Student *t* test; ns, not significant; *p < 0.05, **p < 0.01, ***p < 0.001.

### S1PR5 is critical for iILC2 exit from the intestine while S1PR1 plays a dominant role in regulating iILC2 egress from MLNs

The repositioning of ILC2s from the intestine to the lung is a multistep process: first crossing endothelial wall to enter local lymphatic vessels and trafficking to MLNs, then egressing from MLNs to enter blood stream, and finally reaching the lung and other tissues. We tested whether S1PR1 and S1PR5 modulate different steps of ILC2 migration by using genetic mouse lines. *S1pr5*^−/−^ mice harbored the similar numbers of ILC2s in both lungs and small intestine compared with WT mice (Figure S5A and B), suggesting that *S1pr5* deficiency does not affect ILC2 development or homeostasis. In response to IL-25 treatment, intestinal ILC2s in *S1pr5*^−/−^ mice rapidly proliferated (Figure 5A and B), but failed to migrate to MLNs, blood and lungs (Figure 5C). We also tested whether *S1pr5* deficiency affects ILC2 migration during acute helminth infection. *N. brasiliensis* infection-elicited iILC2s in MLNs and lungs on day 5 post infection were significantly decreased in *S1pr5*^−/−^ compared to WT mice (Figure 5D). These results suggest that S1PR5 is essential for intestinal ILC2s egress from the lamina propria to lymphatics and draining LNs.

**Figure 5.**
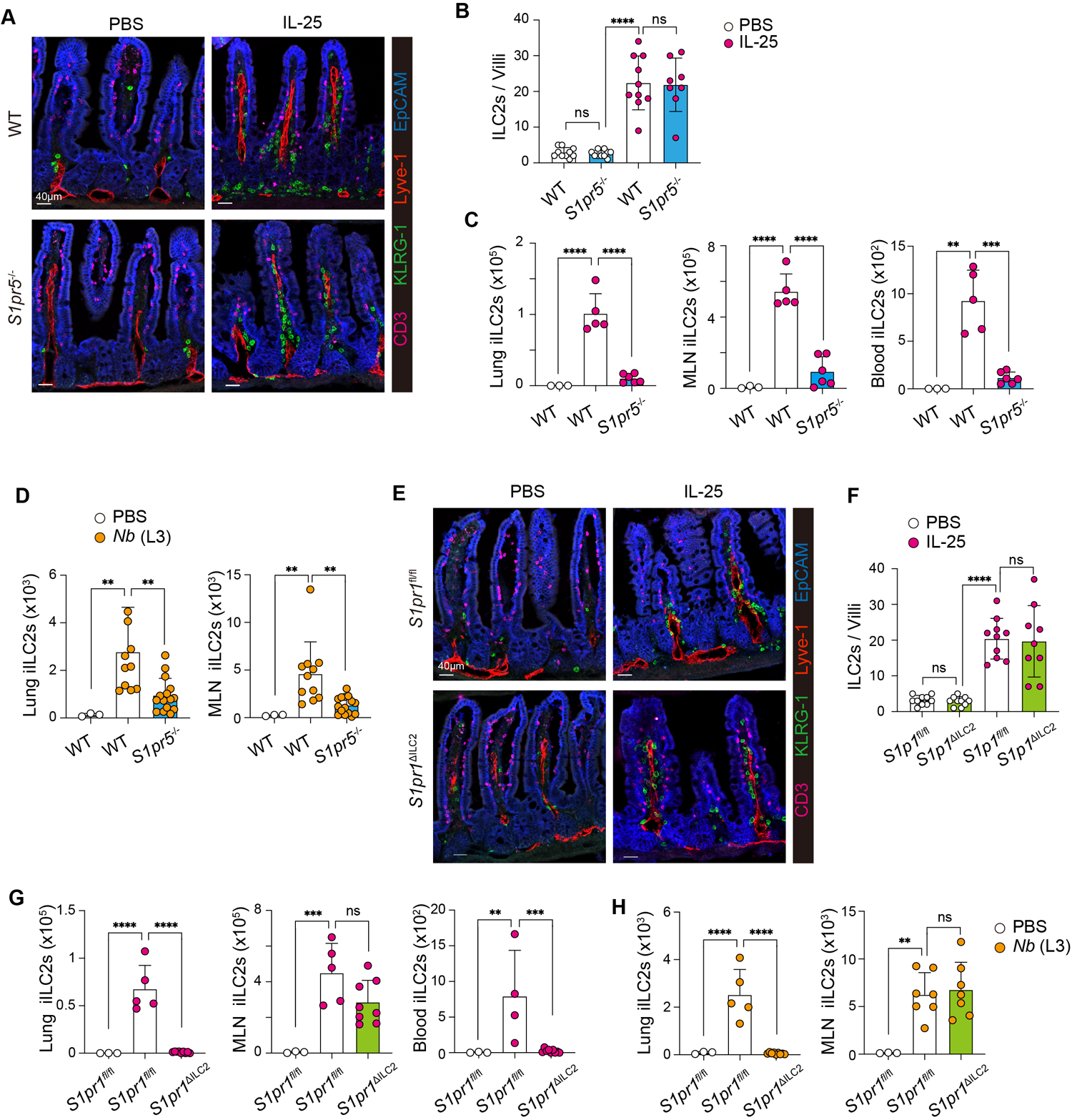
S1PR5 and S1PR1 regulate ILC2 migration in a stage-specific manner. **(A)** Confocal imaging of small intestine from WT or *S1pr5*^−/−^ mice treated with PBS or IL-25 for three days. **(B)** Quantitative analysis of iILC2s per villa in the small intestine of the mice in A. **(C)** FACS analysis of iILC2 cell numbers in lung, MLN and blood of the mice in A. **(D)** Cell numbers of iILC2s in the lung and MLNs of WT or *S1pr5*^−/−^ mice infected with L3 *Nb* for 5 days. **(E)** Confocal imaging of small intestine from *S1pr1*^ΔILC2^ mice or littermate controls treated with PBS or IL-25 for three days. **(F)** Quantitative analysis of iILC2s per villa in the small intestine of the mice in E. **(G)** FACS analysis of iILC2 cell numbers in lung, MLN and blood of the mice in E. **(H)** Cell numbers of iILC2s in the lung and MLNs of *S1pr1*^ΔILC2^ mice or littermate controls infected with L3 *Nb* for 5 days. Data are shown as the mean ± SEM. Student *t* test; ns, not significant; *p < 0.05, **p < 0.01, ***p < 0.001, ****p < 0.0001.

We next investigated the role of S1PR1 during ILC2 migration. We crossed *S1pr1*^fl/fl^ mice with *Klrg1*^Cre^, which was used as an ILC2-specific Cre in our recent publication(*47*). Conditional deletion of *S1pr1* (*S1pr1*^ΔILC2^) did not affect the development or homeostasis of intestinal ILC1, ILC2, ILC3, as well as lung ILC2 (Figure S6A-D). Though a small fraction of T cells express KLRG1(*47*), the total numbers of CD4+ T cells in the lung, spleen and small intestine remained largely unchanged in *S1pr1*^ΔILC2^ mice compared to littermate controls (Figure S6E-G). In response to IL-25 treatment, intestinal ILC2s in *S1pr1*^ΔILC2^ mice underwent rapid proliferation similar to their littermate controls (Figure 5E and F). Strikingly, *S1pr1* deficiency did not impact iILC2 exit from the intestine to MLNs but nearly completely blocked iILC2 migration from MLNs to the blood and lung (Figure 5G). Similar effect was also observed during *N. brasiliensis* infection-induced iILC2 migration (Figure 5H), suggesting that S1PR1 plays an important role in the step of MLN-to-periphery, but not gut-to-MLN, during iILC2 migration. Taken together, these data suggest that S1PR5 and S1PR1 regulate ILC2 migration in a stage-specific manner.

### The tissue-retention marker CD69 inhibits S1PR1 presence on iILC2 cell surface

It was interesting to observe that ILC2s utilize two distinct S1PRs for different migratory steps. RNA-seq results showed that, while migratory iILC2s upregulated S1PR1, they downregulated the tissue-retention molecule CD69 (Figure 6A). In T cells, CD69 binds S1PR1 on cell surface to prompt S1PR1 internalization and degradation(*48*). We hypothesized that CD69-mediated internalization presents S1PR1 protein presence on ILC2 cell surface therefore another S1PR, which is S1PR5, is required for ILC2 exit from tissue. In response to IL-25, gILC2s upregulated *S1pr1* mRNA expression before egressing intestinal tissue (gut iILC2) (Figure 6B). However, S1PR1 protein level on cell surface didn’t increase significantly until iILC2s reached MLNs (Figure 6C and D). By contrast, CD69 remained highly expressed as long as iILC2s stayed in the intestine, but dramatically decreased when they arrived MLNs (Figure 6E and F), indicating a reserve correlation between S1PR1 and CD69 protein levels on iILC2s. Indeed, *Cd69* deficiency led to an increase of S1PR1 protein on gut iILC2s compared to WT cells (Figure 6G and H). *Cd69* deficiency did not affect IL-25-induced iILC2 migration to MLNs (Figure 6I). While S1PR5 antagonist treatment impaired iILC2 migration from the intestine to MLNs in WT mice, it failed to do so in *Cd69*^−/−^ mice (Figure 6I), presumably due to the presence of S1PR1 that can substitute S1PR5 function for S1P sensing. These results suggest that CD69 prevents the presence of S1PR1 protein on iILC2 cell surface, resulting the requirement of another S1P receptor, S1PR5, for iILC2 exit from the intestine.

**Figure 6.**
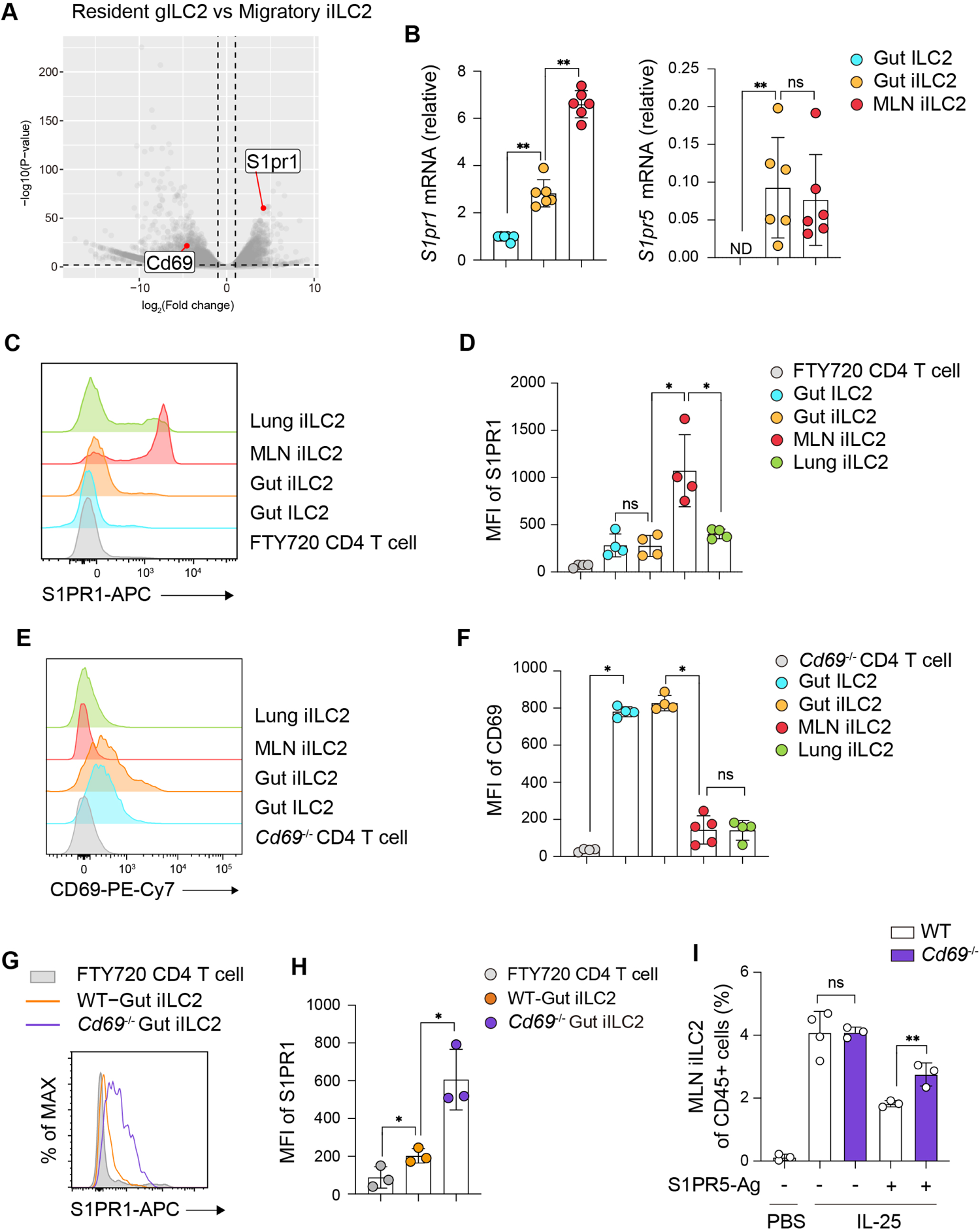
CD69 inhibits the presence of S1PR1 on iILC2 cell surface. **(A)** Volcano plot of differentially expressed genes in RNA-seq between resident gILC2s and migratory MLNs iILC2s. **(B)** RT-qPCR analysis of *S1pr1* and *S1pr5* mRNAs in gILC2s, gut iILC2s and MLN iILC2s. **(C** and **D)** FACS staining and quantification of mean fluorescent intensity (MFI) of S1PR1 on gILC2s, gut iILC2s, MLN iILC2s and lung iILC2s. FTY720-treated CD4 T cells were used as a negative control for S1PR1 staining. **(E** and **F)** FACS staining and quantification of CD69 MFI on indicated cell populations. **(G** and **H)** FACS staining and MFI quantification of S1PR1 on gut iILC2s from WT and *Cd69*^−/−^ mice. **(I)** Frequency of IL-25-elicited MLN iILC2s in WT and *Cd69*^−/−^ mice treated with or without S1PR5 antagonist BIO-027223. Data in **B, D**, **F**, **H,** and **I** are shown as the mean ± SEM. Student *t* test; ns, not significant; *p < 0.05, **p < 0.01.

### The absence of migratory iILC2s leads to prolonged lung tissue damage during *N. brasiliensis* infection

To elucidate the physiological role of S1PR1/S1PR5-mediated ILC2 redistribution, we assessed lung histopathology of WT and *S1pr5*^−/−^ mice on day 8 post *N. brasiliensis* L3 infection. Compared to WT mice, the lungs of *S1pr5*^−/−^ mice exhibited an elevated lacunarity score, which reflected discernible gaps within the alveolar architecture and impaired tissue repair (Figure 7A and B). iILC2 cell numbers in the lung of *S1pr5*^−/−^ mice were significantly decreased compared to WT mice (Figure 7C), while ST2^+^ Th2 cell numbers were similar between these two groups of mice (Figure 7D), suggesting that *S1pr5* deficiency does not affect Th2 recruitment to the lung during *N. brasiliensis* infection. Consistent with the decreased iILC2 cell numbers, *Il5*, *Il13* and *Areg* expression levels were reduced in the lung of *S1pr5*^−/−^ mice compared to WT mice (Figure 7E). Similarly, *S1pr1*^ΔILC2^ mice exhibited elevated tissue lacunarity, decreased iILC2 accumulation and reduced *Il5*, *Il13* and *Areg* expression in the lung during *N. brasiliensis* infection (Figure 7F-J). These two distinct genetic models, which shared a common iILC2-deficient phenotype, showed similar lung pathology, suggesting that the redistributed iILC2s play a critical role in prompting lung tissue repair during *N. brasiliensis* infection.

**Figure 7.**
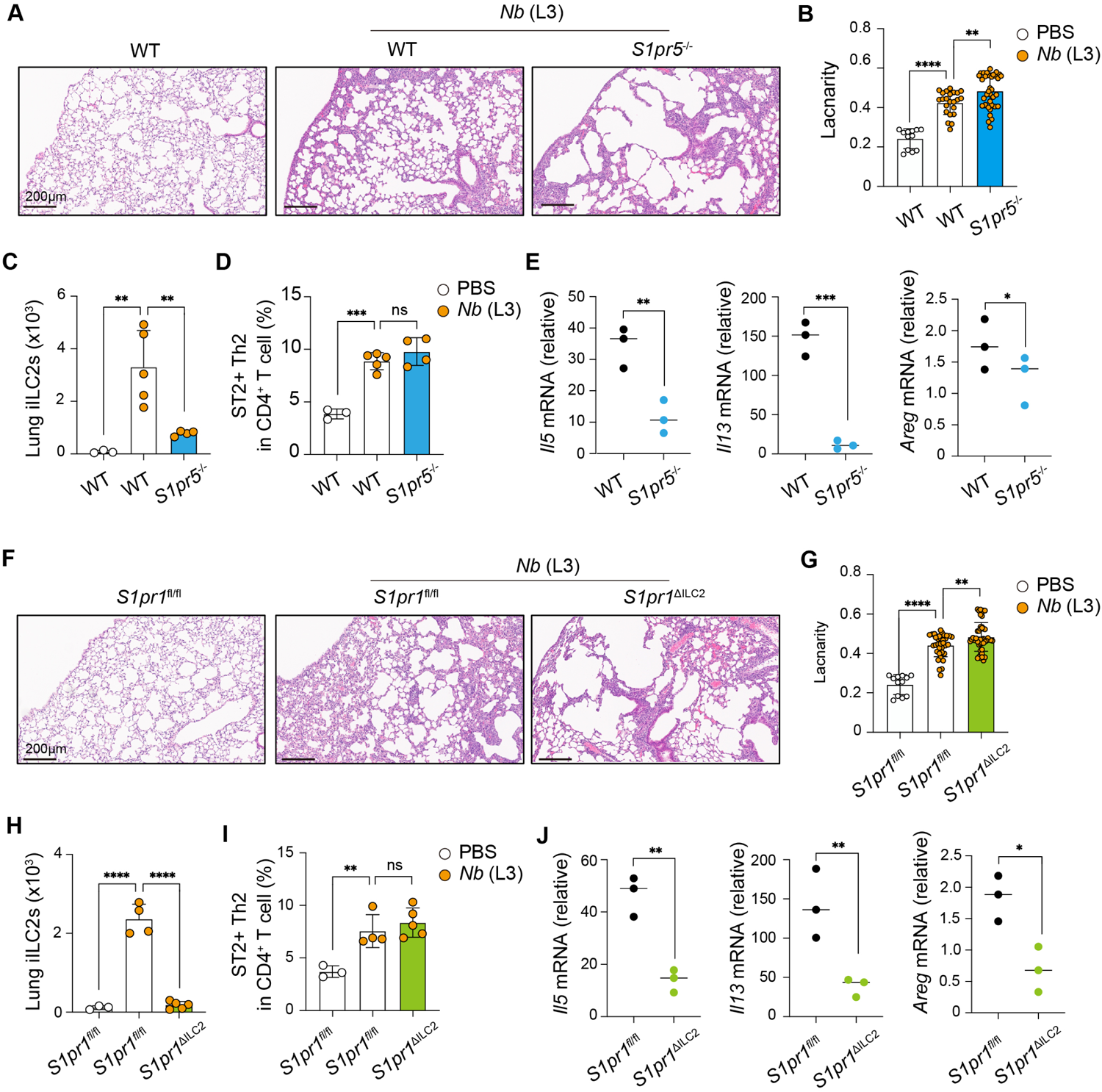
S1PR5/S1PR1-mediated ILC2 redistribution is essential for lung tissue repair. **(A)** Representative H&E staining of lung sections from WT and *S1pr5*^−/−^ mice infected with *Nb* L3 for 8 days. **(B)** Lacunarity score of lung tissues in A. **(C** and **D)** FACS analysis of iILC2s and Th2 cells in the lung of the mice in A. **(E)** RT-qPCR analysis of *Il5*, *Il13* and *Areg* mRNA expression in the lung in A. **(F)** Representative H&E staining of lung sections from *S1pr1*^ΔILC2^ mice or littermate controls infected with *Nb* L3 for 8 days. **(G)** Lacunarity score of lung tissue in F. **(H** and **I)** FACS analysis of iILC2s and Th2 cells in the lung of the mice in F. **(J)** RT-qPCR analysis of *Il5*, *Il13* and *Areg* mRNA expression in the lung in F. Data are shown as the mean ± SEM. Student *t* test; ns, not significant; *p < 0.05, **p < 0.01, ***p < 0.001, ****p < 0.0001.

## Discussion

The recently revised paradigm of ILC residency/migration suggests that, although ILC2s adapt into intestinal tissue during the early development and largely stay as tissue-resident, they are capable of migrating to distal tissue sites during infection(*5, 49, 50*). Here, we illustrate a migratory path of IL-25- or helminth infection-induced iILC2s, which cross endothelium and enter villa lymphatic vessels from lamina propria and then travel through lymphatics to enter blood circulation. It has been proposed that other immune cells such as neutrophils are able to directly re-enter blood vessels from the tissue via reverse TEM(*51*), however, our results of MLN removal and peripheral LNs survey experiments suggested that local lymphatic draining network is the primary route for iILC2 exit from intestinal tissue.

We showed that SLOs including MLNs are redundant for iILC2 interorgan migration and effector cytokine production. This observation is in line with the view that SLOs emerge along with T and B lymphocytes during evolution and serve as a signal hub for adaptive immune reaction(*52*). In WT mice, iILC2s travel through MLNs, providing an opportunity for iILC2s interacting with T cells. ILC2s have been reported to prompt T cell responses via cell-cell interaction mediated by MHC-II or co-stimulatory receptors such as OX40L(*53, 54*). However, whether such a crosstalk between ILC2 and T cells is limited within MLNs or it can also occur in non-lymph tissue sites such as lung and intestine is an intriguing question to be investigated.

We demonstrated a stage-specific requirement of different S1PRs, namely S1PR5 for gut-to-lymph and S1PR1 for MLN-to-circulation. The advantage of using different S1PRs is to allow intestinal ILC2s to sense S1P gradient when one receptor is disrupted. Recent biochemical and structural studies showed that CD69 selectively binds to the transmembrane domain T (TM4) of S1PR1 but not other S1PRs, and acts in cis as a protein agonist of S1PR1, thereby promoting G_i_-dependent S1PR1 internalization(*55*). CD69 is considered as a tissue retention marker and is expressed on other types of tissue resident immune cells such as T_RM_ and γδT cells(*56–58*). S1PR5 is also expressed on T_RM_ and has been suggested to be involved in T_RM_ emigration from peripheral organs(*41*). Therefore, we hypothesize that it is a shared mechanism that CD69-expressing tissue-resident immune cells requires another S1P receptor rather than S1PR5 for egressing from tissues.

It was surprising that IL-25 induced such a drastic change in epigenome landscape of intestinal ILC2s. It has been showed that each ILC lineage possesses unique open chromatin landscapes, and these features are relatively static after ILC activation, revealing the poised status of ILCs prior to stimulation(*39*). Each ILC subset has unique genome-wide distribution patterns of methyl-CpG and 5-hydroxymethylcytosine (5hmC) DNA methylations, which are associated with open chromatin regions, histone modifications and TF-binding sites of corresponding ILC lineage(*59*). The global cellular chromatin architecture of ILC2s remain constant after activation, with rapid remodeling of 3D configuration for type 2 cytokine gene transcription(*60*). We showed here that, in response to IL-25 signal, intestinal ILC2s gains chromatin accessibility in proximity to S1PR genes while the accessibility in Th2 coli remained largely unchanged. How IL-25 induces a selective epigenomic program and how IL-25 signaling impacts DNA methylations and 3D chromatin structure are important questions to be addressed in future studies.

In summary, our study reveals a migratory route of ILC2 repositioning from one tissue to another via lymphatic and vascular circulatory network. While lymph nodes are essential for T cells education and redistribution, they are redundant for the repositioning of ILC2s, which are pre-differentiated and directed by tissue alarmins. This work provides a novel framework for understanding the dynamic repositioning of many tissue-resident immune cells that must navigate the circulation for utility in other tissues. It also sheds new insights to a fundamental paradigm of how S1P modulates both innate and adaptive lymphocytes emigration for lymph or non-lymph organs.

## Methods

### Mice

C57BL/6, *Lta*^−/−^, and *Map3k14*^-/-^, *S1pr1*^fl/fl^ mice were from the Jackson Laboratory. *Klrg1*-cre mice(*61*) were provided by Richard Flavell at Yale University. *Rorc-egfp* reporter mice were provided by Ivaylo Ivanov at Columbia University. *S1pr5*^−/−^ mice(*23*) were provided from Jerold Chun at Sanford Burnham Prebys Medical Discovery institute. *Cd69*^−/−^ mice(*62*) were from Toshinori Nakayama at Chiba University. Mice used for experiments were females between 8-16 weeks of age, unless otherwise specifically indicated. All animal experiments were performed under the approval by the Institute Animal Care and Use Committee of Columbia University. For S1PR agonist treatment, mice were injected intraperitoneally (i.p.) with or without SEW2871 (Cayman Chemical), CYM50358 (Tocris), or BIO-027223 (from Biogen) at the indicated doses combined with IL-25 i.p. injection (200ng/mice) daily for 3 days. For 4-deoxypyridoxine treatment, mice received 30 mg/l 4-deoxypyridoxine (Sigma) combined with 1 g/L sucrose or 1 g/L sucrose alone in the drinking water 3 days before IL-25 injection.

### Mesenteric lymphadenectomy

C57BL/6 were anesthetized and the small intestine, cecum, and MLNs were exteriorized through a 1-cm-wide incision along the abdomen and kept humid with PBS. Mesenteric lymphadenectomy was performed by microdissection along the length of the superior mesenteric artery to the aortic root. After surgery, the small intestine and cecum were reintroduced into the abdomen, the lesion of the abdominal wall stitched with degradable thread, and the outer skin sealed with wound clips. The mice were then subjected to IL-25 i.p injections starting on day 5 post the surgery.

### Nippostrongylus brasiliensis infection

Mice were inoculated subcutaneously with ∼350 third or fifth-stage *N. brasiliensis* larvae (L3 or L5). L5 larvae were prepared as previously described(*33*). Lung, lymph nodes, blood and small intestine tissues were harvested on day 5 or 8 post inoculation for analysis.

### RNA isolation and quantitative PCR

Lung tissue, small intestinal tissue or sorted cells were disrupted in TRIZol Reagent and total RNA was purified according to the manufacturer’s protocol using RNeasy Kits (Qiagen).

Reverse transcription was performed by using PrimeScript™ RT Reagent Kit (Takara). Quantitative PCR was run using SYBR green Master Mix or TaqMan Fast Advanced Master Mix on a QuantStudio 3 (Applied Biosystems) system. GAPDH was used to normalize the RNA content of the samples. The primer sequences for SYBR qPCR are:

GAPDH_For GGGGTCCCAGCTTAGGTTC

GAPDH_Rev TTCACACCGACCTTCACCATT

Areg_For GCTGAGGACAATGCAGGGTAA

Areg_Rev GTGACAACTGGGCATCTGGA

The primer used for TaqMan are GAPDH (ThermoFisher, Mm99999915_g1), *Il5* (ThermoFisher, Mm00439646_m1) and *Il13* (ThermoFisher, Mm00505403_m1).

### Immunofluorescence staining and confocal imaging

The MLN and ileal portion of the small intestine were excised, and small intestine samples were prepared as a “Swiss roll,” then incubated in 4% PFA overnight followed by dehydration in 30% sucrose prior to embedding in OCT freezing media (Sakura Finetek). Twelve-micron sections were cut on a CM3050S cryostat (Leica) and adhered to Superfrost Plus slides (VWR). Frozen sections were blocked in PBS containing 4% Bovine serum albumin (Sigma) followed by staining with antibodies diluted in blocking buffer. The following primary antibodies were used for staining: anti-EpCAM (G8.8, Biolegend), anti-CD3 (17A2, Biolegend), anti-B220 (RA3-6B2, BD Pharmingen) anti-KLRG1 (2F1, BD Biosciences), anti-Lyve-1 (ALY7, eBiosciences). If necessary the secondary antibodies were used at 1:500 dilution at RT for 1 hour. After staining, slides were mounted with Fluormount G (Southern Biotech) and examined on a Nikon Ti Eclipse confocal microscope. Images were analyzed by Image Imaris (Bitplane). The number of iILC2s were counted manually within 10-20 random villus, 25µm x 25µm square of 10-20 random subcapsular area, and 50µm x 50µm of 10-20 random TCZ and BCZ.

### Histology

Lung sections were fixed in 4% formalin buffered for at least 3 days, followed by processing in Tissue-Tek VIP 5 Tissue Processor (Sakura) according to the manufacturer’s instructions. Tissue blocks were paraffin-embedded using Tissue-Tek TEC 5 Embedding Station (Sakura) and sliced into 5-µm sections using a Leica microtome. Tissue sections were then deparaffinized, hydrated and stained with hematoxylin and eosin (H&E). Images were acquired with Aperio AT-2 slide scanner (Leica) and tissue lacunarity was determined by Image-J.

### Antibodies and reagents

The following fluorochrome-conjugated antibodies were used for flow cytometry. Anti-CD3ε (145-2C11), anti-CD5 (53-7.3), anti-CD19 (1D3), anti-B220 (RA3-6B2), anti-CD11b (M1/70), anti-CD11c (N418), anti-NK1.1 (PK136), anti-TCRγδ (eBioGL3), anti-Gr-1 (RB6-8C5), anti-FcεR1 (MAR-1), anti-CD4 (RM4-5), anti-CD8a (53-6.7), anti-CD49b (DX5), anti-TER119 (TER-119), anti-Thy1.2 (30-H12), anti-MHCII (M5/114), anti-CD45.1 (A20), anti-CD45.2 (104), anti-RORγt (AFKJS-9) antibodies were from eBioscience. Anti-Thy1.1 (HIS51) antibody was from Thermo Fisher. Anti-KLRG1 (2F1), Anti-CD69 (H1.2F3), and anti-GATA3 (L50-823) antibodies were from BD Biosciences. Anti-ST2 (DJ8) antibody was from MD Bioproducts. S1PR1 (MAB7089) were from R&D Systems. Recombinant IL-25 were from R&D Systems. The LIVE/DEAD fixable dead cell stain kit was from Life Technologies.

### Flow cytometry and cell sorting

Cells in PBS solution with 3% FBS were blocked with anti-CD16/CD32 (2.4G2, Harlan Laboratories) and then were incubated with fluorochrome-conjugated antibodies with LIVE/DEAD fixable dead cell stain dye. Staining and washing were performed at 4℃. Staining for S1PR1 was done following a previous paper(*63*). Briefly, cells were stained for 90 min with anti-mouse S1PR1 (MAB7089, R&D Systems) on ice, washed twice in buffer, stained for 45 min with anti-rat IgG-biotin F(ab′)2 (2340649, Jackson Immunoresearch) on ice, washed twice in buffer and stained with streptavidin coupled with APC. CountBright Absolute Counting Beads (Life Technologies) were added into cell suspension before analysis. Cells were analyzed on LSR Fortessa flow cytometer (BD Biosciences) and data were analyzed with FlowJo software. ILCs were gated on live CD45^+^ Lineage (Lin)^−^ Thy1^+^, and then further gated on RORγt^−^ NK1.1^+^ as ILC1s, NK1.1^−^ KLRG1^+^ or GATA3^+^ as ILC2s, and RORγt^+^ as ILC3s. iILC2s were gated as live CD45^+^ Lin^−^ Thy1^+/low^ ST2^−^ KLRG1^hi^. Lin antibody cocktail included CD3, CD5, CD19, B220, CD11b, CD11c, TCRγδ, Gr-1, FcεR1, CD8α, DX5 and TER119. CD4^+^ T cells were gated as live CD45^+^ CD3^+^ CD4^+^. For cell sorting, cells were stained and washed in a PBS solution with 10% FBS. Cells were purified on FACS Aria cell sorter (BD Biosciences).

### Isolation leukocytes from lung, LNs, and small intestine

Lung tissues were harvested after perfusion and were disrupted into small pieces, then were digested for 20 min at 37 °C with Liberase TM (Roche) plus DNase I (Roche). Tissue pieces were strained through 40μm cell strainer into single cells followed by treatment with ACK Solution (Life Technologies) for 20s. LN tissues were harvested and were strained into single cells. Blood (100μl) were harvested and were strained into single cells after ACK treatment. Small intestines were harvested and the contents were emptied. Peyer’s patches were removed and then the small intestines were opened longitudinally, cut into small pieces and shaken at 37 °C for 20 min in HBSS media containing 2.5% FBS, 5 mM EDTA and 1 mM DTT for three times to dissociate intraepithelial leukocytes. The remained fragments were washed twice with PBS and then were digested at 37 °C for 20 min in RPMI media containing 5.0% FBS, Collagenase A (Roche), and DNase I (dose?) (Sigma) for three times. The digested tissues were strained to yield a single-cell suspension, which was followed by treatment with ACK Solution. Blood was collected 100µl and were strained into single cells. For detecting S1PR1, we used charcoal-stripped fetal bovine serum instead of FBS.

### Transwell Assay

Transwell migration assays were conducted as previously described(*64*). Following a period of serum deprivation utilizing RPMI 1640 medium supplemented with 0.5% charcoal-depleted fetal bovine serum for a duration of 3 hours at 37°C, iILC2 derived from mesenteric lymph node cells (5×10^4^ cells) were introduced into the upper chambers of 24-well tissue culture inserts (Costar) possessing 5-µm pores fashioned from polycarbonate material. This introduction was facilitated with a volume of 100µl, while the lower chambers were supplied with 600µl of S1P (500 nM) (Sigma Aldrich). In certain experimental iterations, a pre-treatment period of 5 minutes with SEW2871 (Cayman), CYM50358 (TOCRIS), or BIO-027223 (from Biogen) was administered to the cells. The migrated cells recovered from each well were counted by flow cytometry by using Counting Beads as a reference.

### RNA-seq

Total RNA was extracted from sorted cell populations using TRIzol (Invitrogen) and Qiagen miRNeasy Micro Kit according to the manufacturer’s instructions. The quantity and quality of total RNA was assessed by BioAnalyzer (Agilent Technologies). 10ng RNA was used to prepare libraries using Single Cell/Low Input RNA Library Prep Kit for Illumina (New England BioLabs). Libraries with different indexes were pooled and sequenced by an Illumina NovaSeq 6000 as paired-end reads extending 150 bases.

The raw reads were first trimmed and filtered using fastp (version 0.21.0) with default parameters. The trimmed reads were then aligned to the mouse reference genome (GRCm38) using STAR (version 2.5.2a) as implemented in the genomon RNA pipeline (version 2.6.3). The gene counts were obtained from the STAR output using htseq-count (version 2.0.2). The count data were normalized and analyzed for differential expression using DESeq2 (version 1.38.3) with the Wald test and an adjusted p-value cutoff of 0.05. The differentially expressed genes were then used as input for gene set enrichment analysis (GSEA, version 4.3.2) with the GO: Gene Ontology gene sets (MSigDB, version 2023.1). GSEAPreranked analysis was used with the default parameters. We used decoupleR (version 2.5.3), a tool that infers the activity of transcription factors from gene expression data to identify potential regulators of the gene expression changes. We applied the weighted mean method with the CollecTRI gene regulatory network.

### ATAC-seq

ATAC-seq was performed according to published protocol (Buenrostro et al., 2013). 10,000 cells were pelleted and washed with 500ul PBS. Transposition was performed directly on nuclei using 25µl tagmentation reaction mix (Tagment DNA Buffer #15027866, Tagment DNA Enzyme #15027865 from Illumina and Digitonin #G9441 from Promega). After pelleting the nuclei by centrifuging at 500g for 10 min, the pellets were re-suspended in 45µl transposition reaction mix to tag and fragmentalize accessible chromatin. The reaction was incubated at 37°C with shaking at 300rpm for 30min. The fragmentalized DNAs were then purified using MinElute Kit (#28006 Qiagen) and amplified using Next High-Fidelity 2x PCR Master Mix (New England Biolabs #M0541) with Nextera DNA CD Indexes, according to the following program: 72 °C for 5 minutes; 98 °C for 30s; 18 cycles of 98 °C for 10s, 63 °C for 30s, and 72 °C for 1min. Once the libraries were purified using PCR Purification Kit (#28106 Qiagen), they were sequenced for 50 cycles (paired-end reads) on HiSeq2500. ATAC-seq reads were mapped to the mouse genome (mm10 assembly) using WashU Epigenome Browser. Binding Motifs of KLF2, ZEB2, and STAT5a were acquired from JASPAR^57^, and mapped with the FIMO from the MEME Suite in sequence^58^ to see potential binding these targets.

### Statistical analysis

Sample and experiment sizes were determined empirically for sufficient statistical power. No statistical tests were used to predetermine the size of experiments. No samples were excluded specifically from analysis, and no randomization or blinding protocols were used. Data distribution was assumed to be normal, but this was not formally tested. GraphPad Prism software was used for statistical analysis. Differences between two groups were determined by two-tailed unpaired, paired Student’s *t*-test, or One-way ANOVA with multiple comparisons using Prism software (GraphPad). P<0.05 was considered significant. Mean ± SD; ns, none-significant, *P < 0.05, **P < 0.01, ***P < 0.001, ****P < 0.0001.

## Acknowledgments

We thank all members of the Huang lab for discussions and assistance on experiments. We thank R. Flavell, I. Ivanov and T. Nakayama for providing critical mouse lines, Columbia Department of Microbiology and Immunology and the Columbia Stem Cell Initiative core facilities staff for assistance with cell sorting and S. Reiner for critical reading of the manuscript. This work was supported by NIGMS 1R35GM138805 (Y.H.) and HL159675 (J.Q.). T. Ito was also supported by Mandl Fellowship and JSPS Fellowship.

## Author contributions

T.I. designed, performed and interpreted the experiments and drafted the initial manuscript. Y.I., V.G. and W.Z. analyzed and interpreted RNA-Seq and ATAC-Seq data. Y.Z. performed ATAC-Seq assay. R. H. assisted with animal experiments. K. G. supplied BIO-027223 compound. J. C. provided *S1pr5*^−/−^ mice. A. S. and J.F.U. provided *N. brasiliensis.* J.Q. assisted with lung pathology and study design. Y.H. designed the experiments, interpreted the data and finalized the manuscript.

## Competing interests

The authors declare no competing interests.

## Data availability

Genomic sequencing data are available in the Gene Expression Omnibus database (accession number pending).

## Notes

### Competing Interest Statement

The authors have declared no competing interest.

