## Supplemental Data for "ILC2s navigate tissue redistribution during infection using stage-specific S1P receptors"

**Fig. S1**

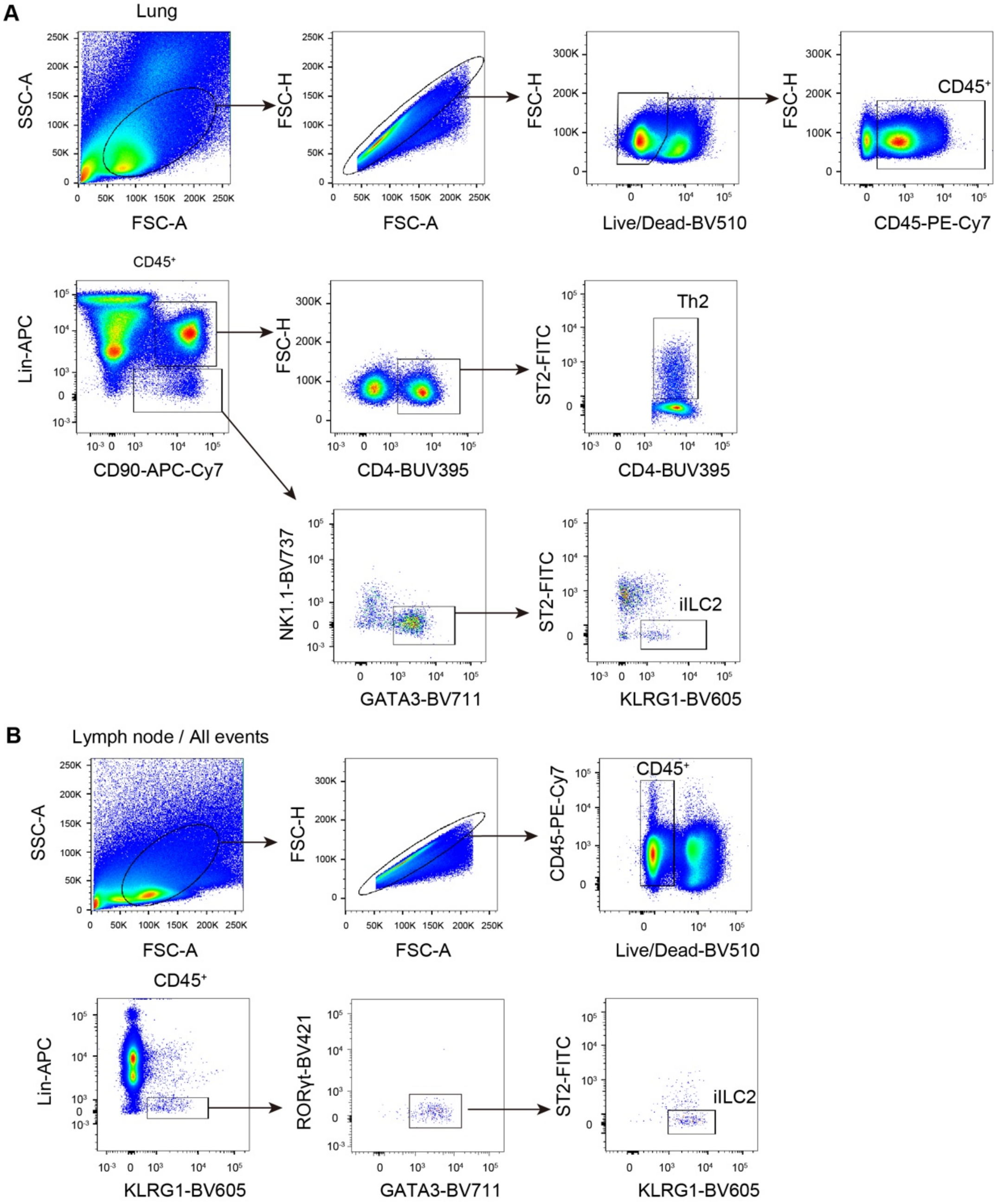


**Supplemental Figure 1. Gating strategy for iILC2.** FACS gating strategy for the phenotypic identification of iILC2 and Th2 in the lung **(A)** and iILC2 in MLNs **(B)**.

Fig. S2

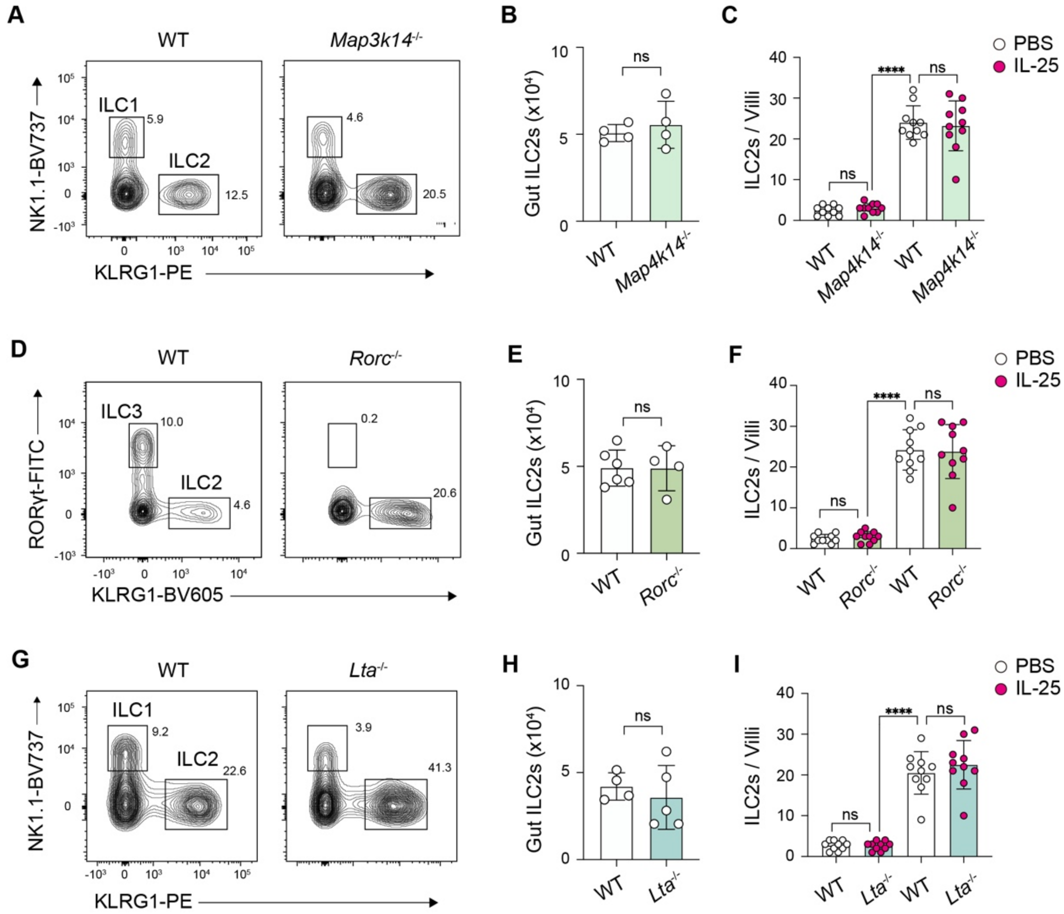


**Supplemental Figure 2. SLO deficiency does not affect ILC2 development and homeostasis.** **(A**, **D**, and **G)** Representative FACS plots of intestine ILC1, ILC2 or ILC3 in three different SLO-deficient mouse lines. ILC2s were gated as live $CD45^+$ $Lin^-$ $Thy1^+$ $NK1.1^-$ (or $ROR\gamma t^-$) $KLRG1^+$. **(B**, **E** and **H)** FACS quantification of intestine ILC2s as in **A**, **D**, and **G**. **(C**, **F** and **I)** Quantitative analysis of confocal imaging of iILC2s per villa in the small intestine. Data are shown as the mean ± SEM. Student *t* test; ns, not significant; *p < 0.05, **p < 0.01, ***p < 0.001, ****p < 0.0001.

Fig. S3

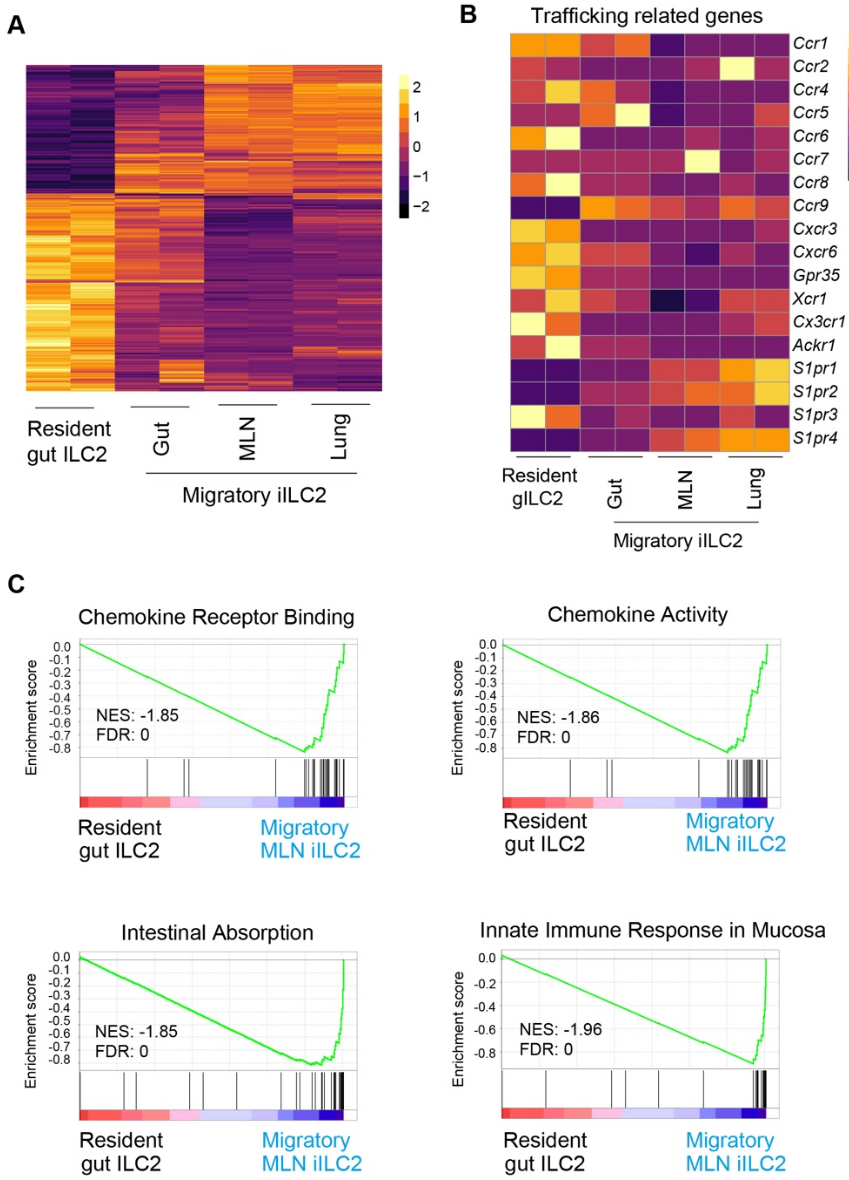


**Supplemental Figure 3. ILC2 transcriptome profile correlates with its migration stage.** **(A)** Heatmap of differentially expressed genes in resident gILC2s and migratory iILC2s located in gut (small intestine), MLNs and lung. **(B)** Heatmap of differentially expressed chemokine-related and S1PR-related genes in resident gILC2s and migratory iILC2s. **(C)** Gene set enrichment analysis (GSEA) of resident gILC2 and migratory MLN iILC2. The normalized enrichment score (NES) and FDR are shown for each plot.

**Fig. S4**

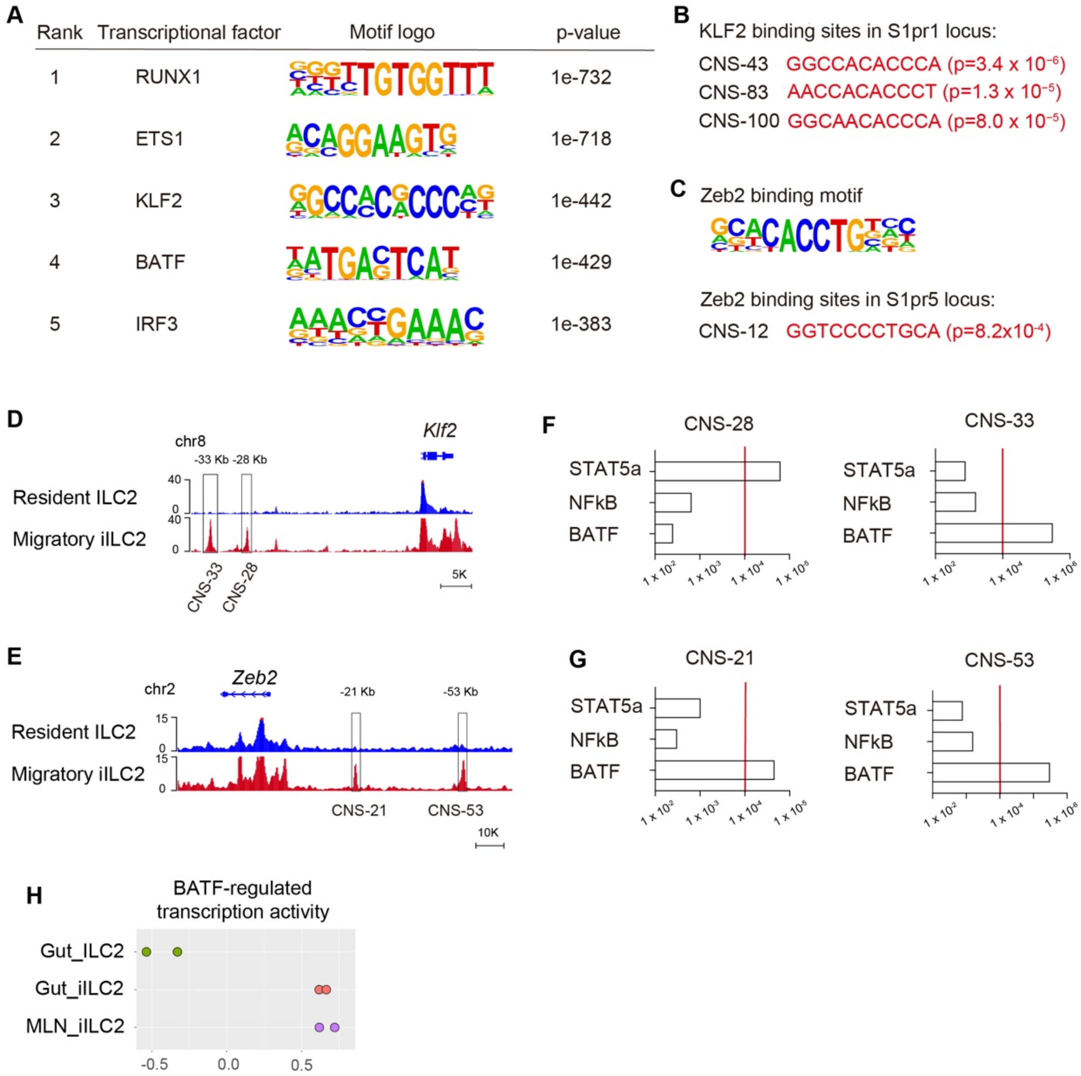


**Supplemental Figure 4. BATF putatively orchestrates the transcriptional program for S1PR1 and S1PR5 expression in iILC2s.** **(A)** Transcription factor binding motif enrichment within the ATAC peaks of migratory iILC2 peaks. The five most significantly overrepresented binding motifs (based on P value) are shown. **(B)** FIMO analysis depicting P values of three KLF2 binding sites in the upstream region of *S1pr1* locus (*CNS-43*, *CNS-83*, and *CNS-100*). The Homer KLF2 motif (shown in A) was used in FIMO analysis. **(C)** Homer ZEB2 motif and FIMO analysis depicting P-values of the ZEB2 binding site in the upstream region of *S1pr5* locus (*CNS-12*). **(D** and **E)** ATAC-seq tracks around *Klf2* locus and *Zeb2* locus in resident gILC2 and migratory iILC2. **(F** and **G)** The enrichment of BATF, STAT5a and NF-κB binding motifs within the accessible peaks close to *Klf2* locus (F) or Zeb2 locus (G). **(H)** Transcriptional activity of BATF in resident gILC2s and migratory iILC2s.

**Fig. S5**

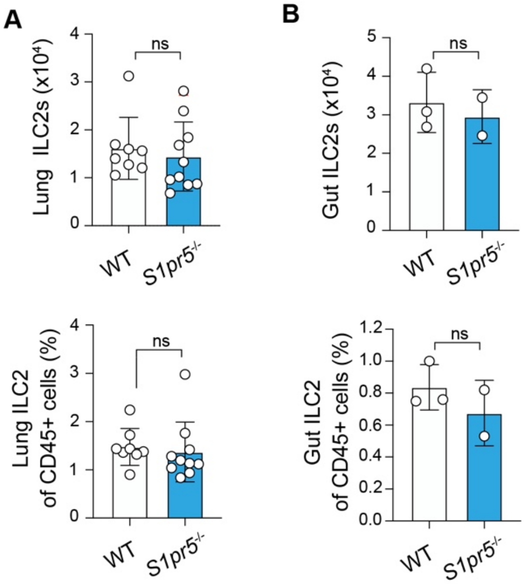


**Supplemental Figure 5. S1PR5 deficiency does not affect ILC2 development or homeostasis. (A** and **B)** FACS analysis of ILC2 number and frequency in the lung and gut of WT and *S1pr5*$^{-/-}$ mice at steady state. ILC2s were gated as live CD45$^{+}$ Lin$^{-}$ Thy1$^{+}$ NK1.1$^{-}$ GATA3$^{+}$. Data are shown as the mean ± SEM. Student *t* test; ns, not significant.

Fig. S6

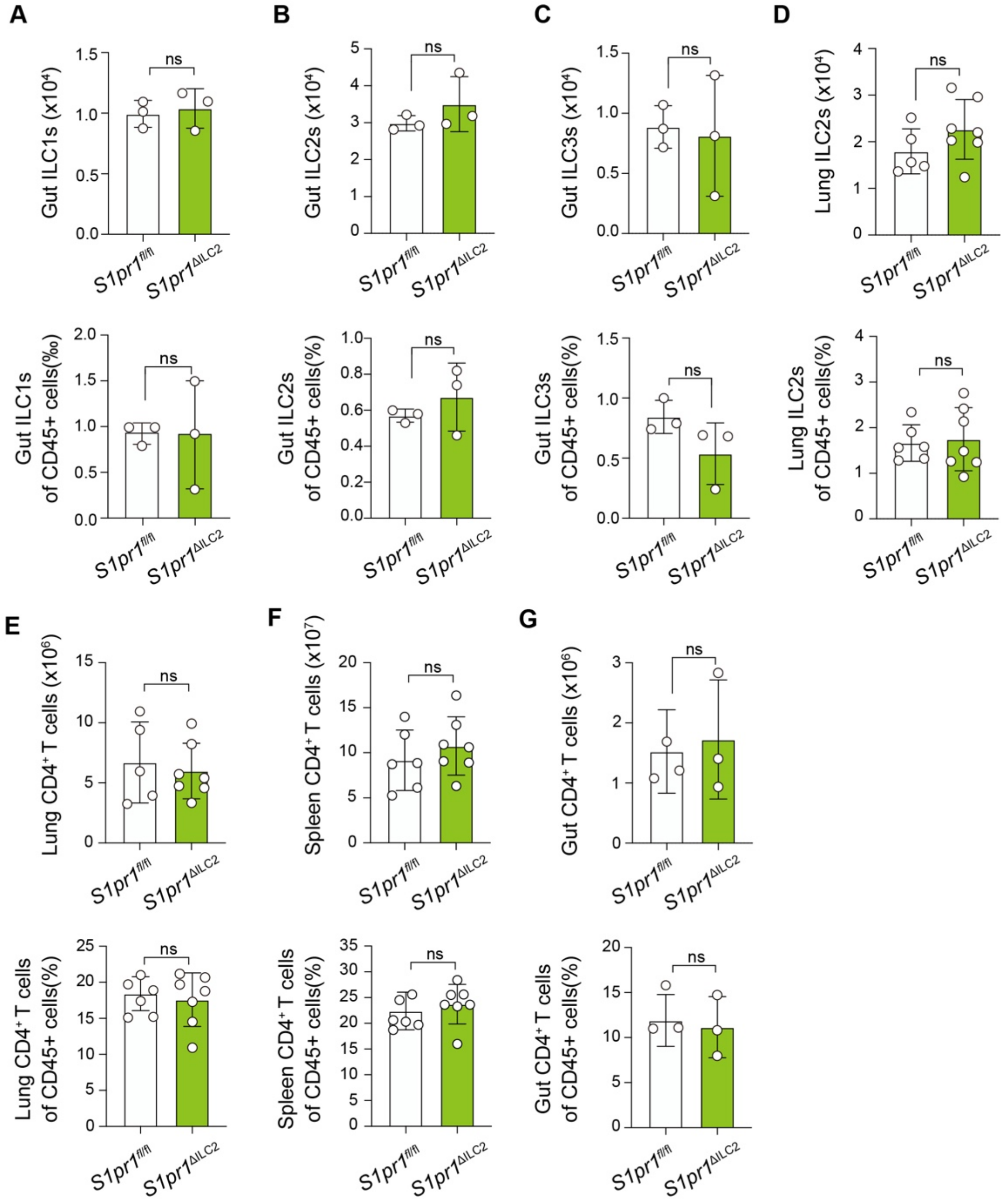


**Supplemental Figure 6. S1PR1 conditional knockout does not affect ILC2 development or homeostasis. (A-C)** Number and frequency of ILC1s, ILC2s and ILC3s in the small intestine of *S1pr1*$^{\Delta ILC2}$ mice and littermate controls. ILC1s were gated as live CD45$^+$ Lin$^-$ Thy1$^+$ NK1.1$^+$ ROR$\gamma$t$^-$; Gut ILC2s were gated as live CD45$^+$ Lin$^-$ Thy1$^+$ NK1.1$^-$ GATA3$^+$; ILC3s were gated as live CD45$^+$ Lin$^-$ Thy1$^+$ NK1.1$^-$ ROR$\gamma$t$^+$. **(D)** Number and frequency of ILC2 in the lung of the mice in A-C. **(E-G)** Number and frequency of CD4$^+$ T cells in lung, spleen and gut of *S1pr1*$^{\Delta ILC2}$ mice and littermate controls. Data are shown as the mean ± SEM. Student *t* test; ns, not significant.
